# Hip osteoarthritis: A novel network analysis of subchondral trabecular bone structures

**DOI:** 10.1101/2022.03.28.486155

**Authors:** Mohsen Dorraki, Dzenita Muratovic, Anahita Fouladzadeh, Johan W Verjans, Andrew Allison, David M Findlay, Derek Abbott

## Abstract

Hip osteoarthritis (HOA) is a degenerative joint disease that leads to the progressive destruction of subchondral bone and cartilage at the hip joint. Development of effective treatments for HOA remains an open problem, primarily due to the lack of knowledge of its pathogenesis and a typically late-stage diagnosis. We describe a novel network analysis methodology for micro-computed tomography (micro-CT) images of human trabecular bone. We explored differences between the trabecular bone microstructure of femoral heads with and without HOA. Large-scale automated extraction of the network formed by trabecular bone revealed significant network properties not previously reported for bone. Profound differences were discovered, particularly in the proximal third of the femoral head, where HOA networks demonstrated elevated numbers of edges, vertices and graph components. When further differentiating healthy joint and HOA networks, the latter showed fewer small-world network properties, due to decreased clustering coefficient and increased characteristic path length. Furthermore, we found that HOA networks had reduced length of edges, indicating the formation of compressed trabecular structures. In order to assess our network approach, we developed a deep learning model for classifying HOA and control cases, and we fed it with two separate inputs: (*i*) micro-CT images of the trabecular bone, and (*ii*) the network extracted from them. The model with plain micro-CT images achieves 74.63% overall accuracy while the trained model with extracted networks attains 96.47% accuracy. We anticipate our findings to be a starting point for a novel description of bone microstructure in HOA, by considering the phenomenon from a graph theory viewpoint.

## INTRODUCTION

Hip osteoarthritis (HOA) is a degenerative hip joint disease that leads to progressive damage of articular cartilage^1^, and structural changes of underlaying the subchondral bone that clinically manifest with overall changes in the (*i*) shape of the femoral head, (*ii*) loss of joint space, (*iii*) frequent severe pain, and loss of joint function^2^. Consequently, HOA is considered as a major cause of disability and loss of life quality^3^, with a prevalence of around 10% for people above 65, where 50% of these cases are symptomatic^4^. With an aging global population, the prevalence of osteoarthritis (OA) is continuing to increase^5^, and is associated with escalating healthcare costs^6^. Thus, optimal management of HOA is of vital importance, which relies on a better understanding of the underlying factors for disease initiation and progression.

It has become clear that events in the subchondral bone under the articular cartilage are intimately involved in the development of OA^7^. In particular, it is well documented that HOA involves pathological changes in the trabecular bone of the femoral head^8,9^. The sequence by which these abnormalities contribute to disease initiation and progression and how they develop, has not been elucidated^10^.

The subchondral bone together with articular cartilage forms an “osteochondral” functional unit with its primary role to maintain joint function. Once synergy between cartilage and subchondral is disrupted, significant structural changes occur in the whole joint. The application of high-resolution imaging approaches such as computed tomography (CT) and magnetic resonance imaging (MRI), in the evaluation of OA patients, has enabled detection of specific OA tissue characteristics in the osteochondral unit and expands the possibilities for diagnosis of disease at an early stage. Of the particular interest are structural changes within subchondral bone. Both animal and human studies indicate that changes of subchondral bone take place early and may precede changes in the articular cartilage and thus it may contribute to not only the initiation but progression of disease^11^.

Subchondral bone immediately beneath cartilage comprises two parts: the subchondral plate and subchondral trabeculae. The subchondral plate is formed by a thin layer of dense bone, from which arise thin trabeculae, forming an intricate and complex network of trabecular bone. Subchondral sclerosis, which encompasses thickening of both the subchondral plate and subchondral trabeculae, is commonly observed in advanced OA and is thus considered as a hallmark of OA^12^. However, it has been reported that in the various stages of OA, different microstructural changes of subchondral bone occur, and it depends on the distance from the articular surface^13–17^. For example, in both humans and animal models of disease during early stages of OA, elevated bone remodelling, subchondral bone plate thinning, and increased porosity were shown to be significantly correlated with cartilage damage^13^. Also during the late stage of OA, appositional bone tissue growth is noted and results in increased subchondral bone apparent density, thickening of the subchondral bone plate, increased bone volume, decrease of trabecular separation, increased trabecular thickness, and change of trabeculae from rod-like into plate-like structures, which associate temporally with articular cartilage thinning and deterioration^10^. Regardless of elevated bone volume density, subchondral bone is less stiff, less mineralized, and less able to withstand repetitive loading^18^.

First, described by Wolff^19^, the bone undergoes dynamic bone remodelling to adapt to the loads, to which it is subjected, and to maintain structural and mechanical integrity. Later, Radin and Rose proposed that an increase in bone density may potentially lead to elevated stiffness that introduces unfavorable stress concentrations resulting in damage to the cartilage^20,21^. More recently, it was suggested that conventional assessments such as thickening of trabeculae, bone volume density, and changes of and thickening of trabeculae structure model index can be insensitive to more subtle changes, and that application of modern image analytical approaches such as individual trabecula segmentation (ITS), are necessary for detection of more early and subtle changes in the properties of the subchondral trabecular bone structure in health and disease^22–24^.

A machine learning approach is proposed to assess the ability of semi-automatically extracted MRI-based radiomic features from tibial subchondral bone to distinguish between knees without and with osteoarthritis. Although the approach is able to classify OA and normal cases, it is not able to explain the link between individual radiomic features and biological processes^25^.

Also, a deep learning approach is used for grading hip osteoarthritis features on radiographs. In a similar way with previous approach, this study used machine learning as a ‘black box’ classifier and it is not able to reveal relation between OA process and visual features in radiographs^26^.

Here, for the first time we propose a novel ‘network’ approach to analyse the trabecular network based on the concepts of graph theory in the human femoral head in subjects with advanced HOA and made comparisons with aged matched control bone with no history of bone disease. Then we develop a machine learning model to classify HOA and control cases. Despite of mentioned previous works, our network approach is able to explain the link between OA biological process and network visual features.

There are several examples of biological networks, such as metabolic networks, protein interaction networks, neural networks, and vascular networks^27–30^. Several complex systems have been investigated recently from a networks viewpoint that link the different elements comprising them^31^. The development of tumour vascular networks has recently been explored via graph theory^32^. Network analysis can qualitatively and quantitatively reveal vital information on the unique characteristics of biological phenomena. The question we explore here is: can trabecular bone microstructure of the femoral head, described as a network graph, visualized by a group of vertices and edges, distinguish between control and HOA bone? Addressing this question may lead to the development of a framework for understanding the progression of the abnormalities in HOA.

We find that a network approach can distinguish with high accuracy trabecular bone of the femoral head with HOA disease from control tissue, despite the wide variability of presentation of trabecular bone between HOA samples. The extraction of the trabecular architecture obtained from micro-CT imaging into a mathematical graph maps the data from a high dimensional space into a low dimensional space. Using a machine learning model, we demonstrate that the low dimensional network representation retains meaningful properties of the original data, near to its intrinsic dimension.

## RESULTS

### Osteoarthritis structural abnormalities assessed by micro-computed tomography

In this study, high-resolution *ex-vivo* imaging using micro-CT provided an opportunity to study the microstructure in the trabecular bone of the femoral head in HOA patients, compared with unaffected hips. Macroscopically, the HOA femoral heads showed considerable variability, qualitatively and in terms of disease severity. The general characteristics, as shown in Fig. 1a, were described on the basis of damage of cartilage (loss of cartilage volume and integrity), and development of subchondral bone sclerosis, osteophytes and bone cysts. A representative micro-CT image of control trabecular bone is shown in Fig. 1b, while 1c shows images of micro-CT slices of an HOA femoral head.

**Fig. 1.**
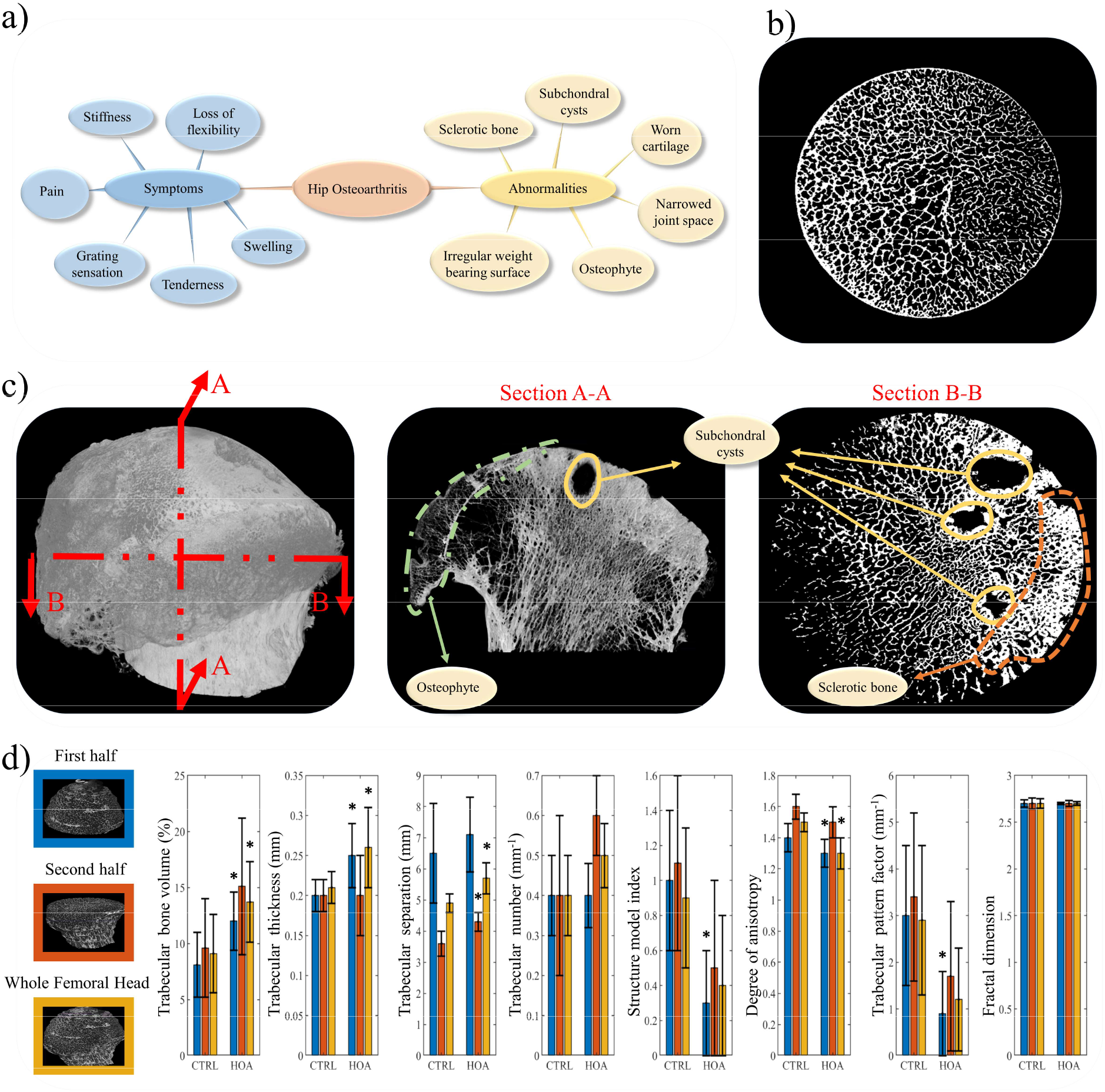
Femoral head abnormalities in HOA. a) HOA symptoms and abnormalities are shown. b) A micro-CT image of trabecular bone in control femoral head without bone related diseases and c) micro-CT images from A-A and B-B sections corresponding to a HOA patient illustrate the abnormalities in trabecular bone. d) An analysis of micro-CT images obtained from a cohort of seven patients with HOA and a cohort of seven CTRLs and seven HOA shows the parameters of trabecular thickness, trabecular bone volume, structural model index, trabecular separation, trabecular number, degree of anisotropy, trabecular bone pattern factor and fractal dimension. Means and standard deviations are shown on the bars for the first half (blue), second half (orange), and whole femoral head (yellow). Here, * indicates statistical significance (*p* < 0.05) between CTRL and HOA. The comparison among the group were performed using unpaired *t*-test.

The femoral heads obtained from subjects undergoing hip replacement surgery and from control (CTRL) individuals without a history of bone disease were scanned using micro-CT (Skyscan 1076, Kontic, Belgium, isotropic resolution = 20.5 μm). After acquiring the micro-CT images, the numerical (trabecular number, bone volume fraction, thickness and separation) and topographic (degree of anisotropy, structural model index, and trabecular pattern factor) properties of subchondral bone microstructure were assessed. Using NRecon software, v2.0.4.2, data were analysed separately for three volumes of interest: whole sample (total of 1950 images = 40 mm), first half of the specimen (images 0 – 975 = 20 mm), the proximal portion of the femur, which articulates with the acetabulum of the pelvis, and the second half (images 976 – 1950, = 20 mm), the distal portion, towards femoral neck.

The results presented in Fig. 1d for the first half (blue bars), the second half (orange bars), and whole femoral head (yellow bars) indicate that HOA trabecular bone had increased bone volume, due mainly to an increased trabecular thickness compared to controls. A similar finding was seen by comparing the trabecular thickness and separation, comparing whole volumes of the femoral head. Trabecular separation is described as the thickness of the spaces measured after image binarization and its increase in HOA samples is consistent with presence of a number of bone cysts present in volume of interest. By contrast, the HOA trabecular bone showed decreased topographic properties, such as structure model index, degree of anisotropy, and trabecular pattern factor, compared to controls. Structure model index shows the relative prevalence of plates and rods. Isotropy is an indication of 3D symmetry or the presence or absence of preferential alignment of structures through a specific directional axis. Trabecular pattern factor is a measure of connectivity, where a highly disconnected trabecular structure is indicated with a higher trabecular pattern factor. The results did not show any significant difference for fractal dimension between HOA and CTRL cases. Fractal dimension is a measure of surface complexity of an object, that quantifies how an object’s surface fills space.

Although these parameters provide useful information about the whole femoral head, they are volume-based parameters and do not shed light on the effects of HOA at different depths. In addition, the large standard deviations in Fig. 1d indicate the presence of uncertainty as the data points are far from the mean. Therefore, defining a more accurate framework dependent that is able to reveal the differences caused by HOA across the depth of the sample, is desirable.

### Hip osteoarthritis: A network analysis

Next, we examined the trabecular networks imaged as micro-CT image slices, obtained from the HOA and CTRL femoral heads. A section of hip bone indicating cartilage, subchondral bone and trabecular bone from proximal (top) to distal (bottom) is shown in Fig. 2a. As described previously, we found that HOA trabecular bone was structurally disordered; this was mainly due to increased trabecular thickness. We analysed the trabecular bone in HOA and CTRL groups as a set of networks and investigated the behaviour of networks as a function of bone depth. As an example, four sections at various depths (5, 15, 25 and 35 mm) are shown in Fig. 2b, and the corresponding HOA and CTRL micro-CT images are shown in Fig. 2c, d. It may be seen that the CTRL trabecular structure appears more uniformly distributed and that the network pattern in both cases changes as a function of depth.

**Fig. 2.**
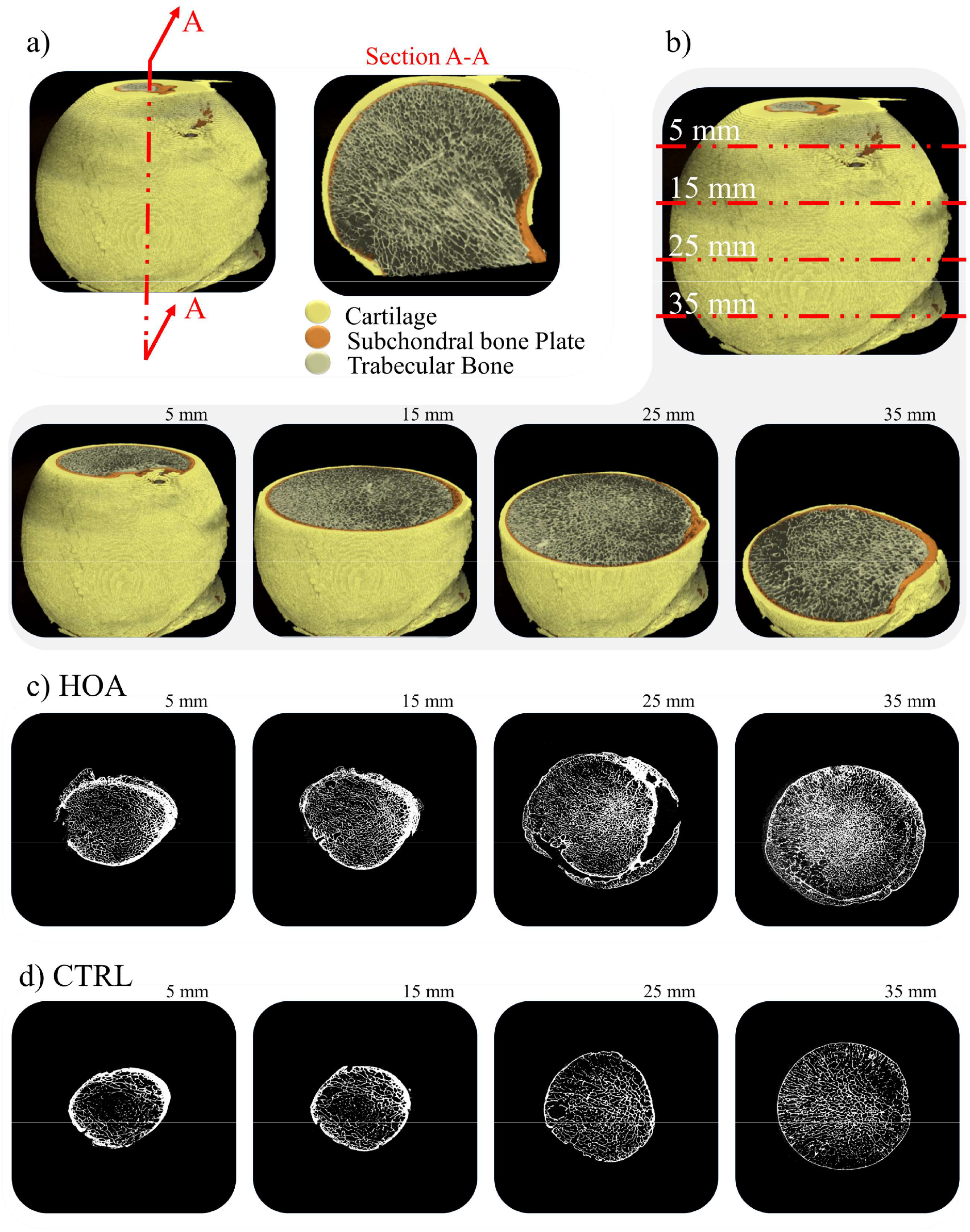
Trabecular bone network changes as a function of depth. a) A section of hip bone showing cartilage, subchondral bone and trabecular bone. b) Four sections at various depths, 5, 15, 25 and 35 mm, are considered, and the corresponding c) HOA and d) CTRL micro-CT images are shown. Although the CTRL trabecular structure appears smoother, the network pattern in both cases changes as a function of depth.

To investigate the trabecular bone network in HOA samples, we developed customised image processing software using MATLAB that is able to extract 2D networks in micro-CT images and display the vertices, edges and several graph parameters in the networks. As an example, in Fig. 3a, a micro-CT image is shown on the left side, and the extracted edges (red lines), vertices (yellow nodes), and branches (green nodes) are illustrated on the right. To further understand the network behaviours, the networks for a HOA and a CTRL case were ‘regraphed’, and the networks were visualized using circular layouts, as shown in Fig. 3b. It can be seen that the entire number of vertices and edges relating to the HOA network is greater than for CTRL, indicating the formation of compressed trabecular structures.

**Fig. 3.**
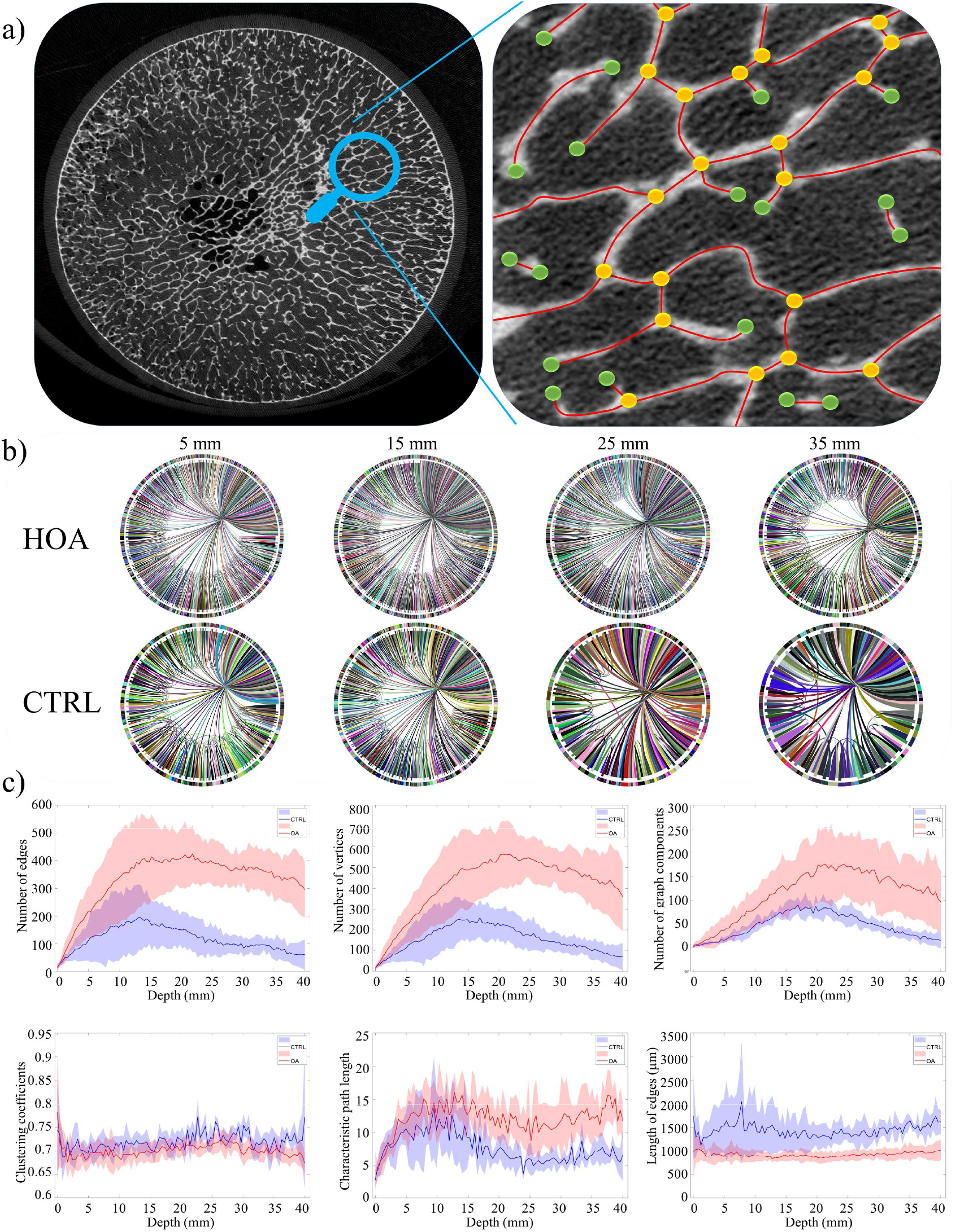
Hip osteoarthritis network analysis. a) An example of identifying edges and vertices from micro-CT images showing networks in HOA and CTRL groups. b) Circular layout of HOA and CTRL corresponding to an HOA and a CTRL case (shown previously in Fig. 2c, d) demonstrating a reducing number of vertices and edges at various depths. c) We analysed micro-CT images obtained from seven HOA (red) and seven CTRL (blue) cases and calculated mean (shown with solid lines) and standard deviation (shaded areas). The corresponding number of vertices, number of edges, number of graph components, clustering coefficients, characteristic path length, and length of edges show the evolution of the bone network in HOA and CTRL cases as a function of depth.

Complex biological networks are often characterized by nonlinearly interacting parameters. To study this intricate connectivity, we further investigated several network parameters, in particular number of vertices, number of edges, number of graph components, clustering coefficients, characteristic path length, and length of edges in Fig. 3c. To achieve this, we analysed micro-CT images from HOA (red) and CTRL (blue) groups—solid lines and shading indicate mean and standard deviation, respectively.

For the micro-CT slices close to the top of the femoral head, the number of vertices and edges grows faster for HOA cases; however, when they reach the maximum point at the first third (top) of the femoral head, both HOA and CTRL decline gradually at the similar rate. These observations show that although HOA trabecular bone has a greater number of vertices and edges, the network in CTRL is more connected, as HOA possesses more graph components than CTRL. Notable also, was a similar trend in the clustering coefficients in HOA and CTRL; however, it may be seen that there is a minor increase in the HOA clustering coefficients. The characteristic path length in HOA is greater that CTRL and both show a fluctuating pattern. The shorter length of edges in HOA indicates the impact of the abnormalities caused by sclerotic changes, increased anisotropy and elevated plate/rod ratio.

The method given here for presenting HOA and CTRL micro-CT images via a network framework can be integrated potentially into machine learning models for HOA diagnosis. As future intelligent disease diagnosis relies on optimising machine learning methods and focusing on data-centric approaches, our network representation shows promise as an effective feature for perform machine learning classification.

### Network properties: Effective feature for machine learning models

To evaluate the outcome of network analysis, we employed a machine learning model for classifying HOA and CTRL. We developed a deep convolutional neural network (CNN) and fed it with two separate inputs (*i*) micro-CT images of the trabecular bone, and (*ii*) the network extracted from the images.

With the advent and progress of artificial intelligence, researchers have been attempting to employ deep neural networks as a novel approach for diagnosis based on clinical and medical data. As machine learning is evidence-based and can analyse problems in an unbiased way, it can be helpful for making objective diagnosis from biomarkers or clinical data^33,34^. Among the deep learning architectures, CNN is particularly suitable for medical image classification due to its ability to take advantage of natural image properties, such as shared weights, local connections, pooling and the employ of several layers to perform preferred analysis^31^, and also it is able to perform the data processing tasks in the form of multiple layers.

Several CNN architectures are used widely for medical image diagnosis, including mitosis detection in breast cancer^35^, deformable registration of MR brain images^36^, and classification of skin cancer^37^. Recently the advent of machine learning systems has attracted the attention of OA researchers. A classification approach based on a probabilistic neural network (PNN) classifier was proposed for the characterisation of hips from pelvic radiograph images as OA or normal^38^. Textural features in this study were extracted from X-ray images. The model accomplishes a high classification rate; however, a limitation is that it lacks automated performance, as it needs a manual procedure via a graphics cursor for recognising regions of interest (ROI) in the X-ray images.

We designed a multiple layer CNN architecture and trained it on 90% of our dataset. Here, Fig. 4 shows the architecture of the proposed CNN deep network. To evaluate the network analysis carried out in previous section, we trained the model first with the micro-CT images, and then assessed the classification output. Then, in a separate procedure, we fed the model with the extracted networks. The database contains *i*) 360 OA and 360 control micro-CT images that are composed of human expert-labelled images, and *ii*) the corresponding networks. We divided the dataset under analysis into training (90% of images) and test (10% of images) sets. After training the deep CNN, we validated the performance of the approach using the test data. The CNN trained on plain micro-CT images achieved 74.63% overall accuracy while the CNN trained on extracted networks attained 96.47% accuracy on a subset of the test set. Here, Fig. 4 summarises our results for CNN model during both experiments.

**Fig. 4.**
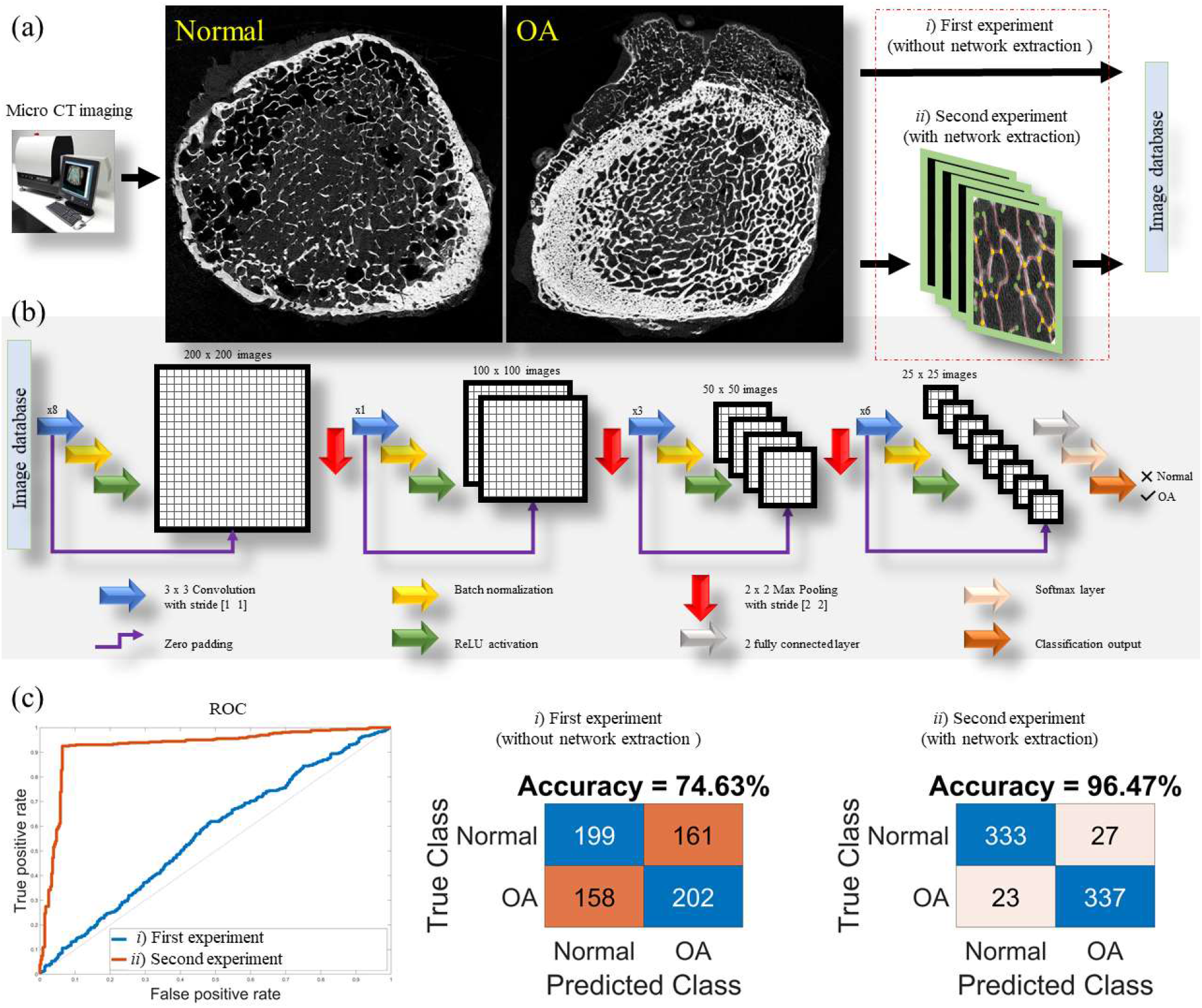
Machine learning model overview. (a) The system includes micro CT imaging and a pipeline of preprocessing techniques fed separately with raw images and extracted networks. (b) Schematic of the CNN operation. It comprises four blocks, each of which are made of a 3 × 3 convolution layer with stride [1 1], a batch normalization and finally a ReLU function. Also, three max pooling layer with stride [2 2] reduce the size of images is used in this architecture. It is followed by two fully connected layers, a Softmax layer and a classification output. (c) Here, a ROC curve for the approach is shown in blue (without network extraction) and red (with network extraction) lines, where the second experiment outperforms the first at the task of classification. In addition, the confusion matrices are illustrated for both experiments.

## DISCUSSION

In this study, we investigated a novel approach for analysing the microstructure of the trabecular bone in HOA, compared with healthy control bone, from a graph theory perspective. By applying our software to high-resolution micro-CT images to analyse the entire trabecular microstructure at various depths, we discovered significant differences in the network configuration in HOA compared with controls that were not apparent using conventional outcome measures.

Imaging approaches such as MRI and radiography have been used widely to diagnose hip osteoarthritis^39^. Unlike plain radiographs, CT can quantify femoral and acetabular forms and evaluates anatomic relationships independently of patient position, and through several postprocessing methods is able to render a 3D visualization of the hip that simplifies observation of bone morphological features. The CT scan provides an accurate assessment^40^ of the bone anatomy of the hip with a high accuracy of 1° to 4.5°. Despite all these advances in imaging technology, how these OA abnormalities change as a function of depth into trabecular bone was unknown until now.

The development of high-resolution imaging, along with the advent of powerful computational approaches, has attracted recently the attention of HOA researchers. The imaging technologies including peripheral quantitative computed tomography (pQCT), magnetic resonance imaging (MRI), dual-energy X-ray absorptiometry (DXA), and micro-CT have revealed main evidence that reveals the significant role of trabecular bone in the pathogenesis of HOA. The high-resolution micro-CT used for the present *ex-vivo* study exceeds the resolution provided by current clinical imaging. However, high-resolution pQCT is rapidly approaching a resolution sufficient for a version of the analysis described here, suggesting that detection of bone abnormalities at early stages may soon be possible, offering the potential for early identification and treatment of these abnormalities. This in turn may potentially lead to attenuation of cartilage damage and the prevention of OA progression^41–44^.

To understand more fully the changes that take place in the subchondral bone in HOA, we assessed the network properties within bone microstructure of femoral head of HOA cases and CTRL bone. Our results show important changes in trabecular bone parameters, with increases in mean trabecular thickness, bone volume, and trabecular separation in HOA, and a concomitant decrease in degree of anisotropy, trabecular pattern factor and structure model index. The novel feature of this study is the inclusion of network parameters. Our results show that the most significant network changes in HOA such as the number of edges, vertices and graph components appear in first third (most proximal) of the femoral head (from top to 1300 μm), and after that both HOA and CTRL networks show similar behaviour. This suggests that subchondral bone changes relate to the OA disease and do not pre-exist the disease. We suggest that these changes and abnormalities in trabecular bone proximal to the diseased joint will open up new avenues for designing artificial intelligence (AI) approaches for HOA diagnosis and classification.

Our analyses reveal that HOA networks are characterised by reduced length of edges, indicating the compressed trabecular structures in patients with HOA. Also, HOA networks show elevated characteristic path lengths and reduced clustering coefficients, therefore lowering the small world network characteristics as they evolve. A network possesses small world properties where its characteristic path length is relatively low, and where its clustering coefficient is relatively high^45^. A high clustering coefficient occurs with highly connected groups, while a short mean path length occurs with rapid information spread^46–48^.

Additionally, we demonstrated the effectiveness of deep learning for hip osteoarthritis classification via plain micro-CT images and the extracted networks. Using a CNN trained on both datasets, we have shown that employing our network approach for feature extraction can significantly increase the classification accuracy. Although we highlight that clinical hip osteoarthritis diagnosis is based on several factors beyond the morphological features in CT scan images, the ability to diagnose OA with high accuracy has the potential to enable earlier intervention and to expand access to vital medical care.

We anticipate our findings to be a starting point for a more sophisticated description of bone microstructure in HOA, by considering the phenomenon from a graph theory viewpoint. Network-based quantitative measures potentially open new avenues for assessing trabecular bone microstructure at other skeletal sites and in other skeletal disease states.

Because OA aetiology is likely multifactorial, and comprises multiple phenotypes, in this study only subjects with primary radiographic and symptomatic OA were included to ensure that detected changes in subchondral trabecular bone are related to this disease. Several diseases such as: osteoarthritis, rheumatoid arthritis, osteoporosis, and psoriatic arthritis are characterized with severe structural changes in subchondral bone. Note that the subchondral bone changes in all these diseases are elicited mainly due to altered bone turnover but their manifestations are various from bone erosion to bone sclerosis^49^.

One of the limitations in this study is that the model did not assess other abnormalities that may co-exist with hip osteoarthritis. However, in this study our focus was to investigate and document if a machine learning method can be used to inform us about bone changes beyond increase/decrease of bone trabecular volume and bone mineral density in the whole femoral head rather than with small volumes of interest that are often used in animal and/or human studies. Therefore, this method may potentially be adapted and used in future clinical HOA studies to benefit medical practice.

A second limitation of our study regards the small number of samples. Although the number of samples in this study was sufficient for statistical investigation and building network analysis, more examples are needed to prove the generality of our findings and to increase the accuracy of our machine learning model. This study is also a cross-sectional study and could not evaluate how the bone structure changes from early stage to late-stage HOA disease. In general, it is challenging to analyse bone microstructure in early stage HOA patients by micro-CT, since the bone samples are not accessible and/or cadaveric tissue with early HOA changes is not easily available. Thus, it will eventually be of clinical interest to investigate a larger cohort longitudinally to determine if our method might play a useful role in assessment of bone across disease severity and/or may help to differentiate osteoarthritis from other diseases of the joint.

In addition, micro-CT cannot be utilized in clinical investigations. However, clinical CT resolution is rapidly improving and *in-vivo* analyses of bone microstructure are becoming increasingly possible as this imaging technology progressively advances.

Moreover, parameters describing subchondral trabecular bone structure in HOA (bone volume fraction, trabecular bone pattern factor, trabecular number, trabecular thickness, trabecular separation, and structure model index) that are often described by micro-CT are already adapted and evaluated *in-vivo* using multi-detector row CT^50–52^. More recently it was elegantly demonstrated that use of clinical CT is a new and promising imaging tool in clinical assessment of both hip and knee OA^53–55^. Thus, we believe that our method will further enrich current knowledge about subchondral bone involvement in pathogenesis of HOA.

## MATERIALS AND METHODS

### Human material

Seven femoral heads (4 males and 3 females aged 71.7 ± 14.6) were collected from patients who have undergone total joint replacement for HOA at the Royal Adelaide Hospital and seven cadaveric femoral heads (5 females and 2 males aged 65.8 ± 15) accessed through the SA Tissue Bank, SA Pathology, Royal Adelaide Hospital Mortuary.

Inclusion criteria for HOA subjects were radiographic HOA with severe symptomatic disabilities including limited mobility and severe pain. Inclusion criteria for non-disease subjects were no evidence of radiographic HOA or joint pain in medical history. Exclusion criteria for both groups: osteoporosis, rheumatoid arthritis, metabolic bone disease, history of malignancy, and medication that may have affected bone turnover. Written consent was obtained for all subjects and the study received prior approval from the Human Research Ethics Committee.

### Data collection

To collect the data related to the microstructure of the trabecular bone, whole femoral head samples were scanned using a micro-CT scanner (Skyscan 1276, Skyscan-Bruker, Kontich, Belgium). Scanner settings: 20.5 μm isotropic pixel size, source voltage 100 kVp, current 200 μA, rotation step 0.4°, 180° rotation, exposure time 700 mm, 3 frames averaging.

For individual femoral head this generated 3,305 X-ray projection images (826 projections per step), image resolution was 3,872 × 3,872 pixels (79,453 × 79,453 μm) in size with depth resolution of 20 μm, in 16-bit Tiff format, producing a total dataset of 24.3 GB, scan duration approximately 4 h.

To insure both adjusting for clustering (multiple regions of interest) and impact of age and sex, data were analysed in a series of separate linear mixed effects regression models to control the confounding of age and sex as independent variables.

### Statistical analysis

The Shapiro-Wilk test was employed to test normality of the data distribution. Differences between groups (Control vs. HOA) were described using the unpaired *t*-test. We performed the statistical analyses via GraphPad Prism software (Version 9.2.0 for MacOS). Data is reported as the mean ± standard deviation (SD). The critical value of *p* < 0.05 was chosen for significance.

### Deep learning model

Our convolutional neural network (CNN) model shown in Fig. 4b contains four blocks, each of which comprise a 3 × 3 convolution layer with stride [1 1], a batch normalization layer and finally a ReLU function. In addition, three max pooling reduces the size of images, and the network followed by two fully connected layers, a Softmax layer and finally a classification output.

The layers where filters are applied to the original image are called convolutional layers. Batch normalization is applied to the output of the previous layers allowing every layer of the network to learn more independently. It can be considered as regularisation to avoid overfitting of the model. We used a ReLU activation function to control the output. Note that ReLU is a linear function that will output the input directly in case it is greater than zero, otherwise, it forces a zero output.

In Fig. 4c, a receiver-operator characteristic (ROC) curve and confusion matrices were obtained as model metrics. A ROC curve is a graph illustrating the performance of a classification model at all classification thresholds. The confusion matrix is a table with two rows and two columns that reports FN, TP, TN, and FP.

Here, FN represents false negative, an incorrect prediction in the negative class; TP represents true positive, a correct prediction in the positive class; TN is true negative, a correct prediction in the negative class; and FP represents false positive, an incorrect prediction in the positive class.

## ACKNOWLEDGEMENTS

Funding sources include a project grant funded by the National Health and Medical Research Council of Australia. The authors wish to acknowledge support from Adelaide Microscopy at the University of Adelaide and Dr Agatha Labrinidis for assisting with micro-CT scanning protocol.

## AUTHOR CONTRIBUTIONS

M.D. developed the network analysis software, performed the computational experiments, and built the machine learning model. D.M. collected the data. M.D. and D.M. wrote the paper. A.F., A.A., and J.W.V. assisted with the computational experiments. D.A. and D.M.F. conceived and supervised the study. All authors proof read paper and checked the analysis.

